# The Evolution of Small-RNA-Mediated Silencing of an Invading Transposable Element

**DOI:** 10.1101/136580

**Authors:** Erin S. Kelleher, Ricardo B. R. Azevedo, Yichen Zheng

**Author notes:** **Associate Editor:** TBD.

## Abstract

Transposable elements (TEs) are genomic parasites that impose fitness costs on their hosts by producing deleterious mutations and disrupting gametogenesis. Host genomes avoid these costs by regulating TE activity, particularly in germline cells where new insertions are heritable and TEs are exceptionally active. However, the capacity of different TE-associated fitness costs to select for repression in the host, and the role of selection in the evolution of TE regulation more generally, remain controversial. In this study, we use forward, individual-based simulations to examine the evolution of small-RNA-mediated TE regulation, a conserved mechanism for TE repression that is employed by both prokaryotes and eukaryotes. To design and parameterize a biologically realistic model, we drew on an extensive survey of empirical studies of the transposition and regulation of *P*-element DNA transposons in *Drosophila melanogaster*. We observed that even under conservative assumptions, where small-RNA-mediated regulation reduces transposition only, repression evolves rapidly and adaptively after the genome is invaded by a new TE. We further show that the spread of repressor alleles is greatly enhanced by two additional TE-imposed fitness costs: dysgenic sterility and ectopic recombination. Finally, we demonstrate that the mutation rate to repression (*i.e.*, the size of the mutational target) is a critical parameter that influences both the evolutionary trajectory of host repression and the associated proliferation of TEs after invasion. Our findings suggest that adaptive evolution of TE regulation may be stronger and more prevalent than previously appreciated, and provide a framework for evaluating empirical data.

## Introduction

Transposable elements (TEs) are intragenomic parasites. Although some TE families or individual TE insertions have been “domesticated” to perform important functions (Kunarso et al., 2010; Lynch et al., 2011; Silva-Sousa et al., 2012) their presence and mobilization generally reduces the fitness of their host. TEs introduce deleterious mutations by causing DNA damage during mobilization and by inserting into functional sequences (Dupuy et al., 2001; Spradling et al., 1999), or disrupting the expression of adjacent genes by inducing heterochromatin formation (Hollister and Gaut, 2009; Lee, 2015; Lee and Karpen, 2017; Sentmanat and Elgin, 2012). Ectopic recombination events between TE insertions in different genomic locations further produce large structural rearrangements, duplications, and deletions, which are overwhelmingly deleterious (reviewed in Hedges and Deininger, 2007). In addition to these mutational impacts, TE activity itself can have drastic fitness consequences by causing dysgenic sterility, a condition in which germline DNA damage prohibits the production of viable gametes (Bingham et al., 1982; Josefsson et al., 2006; Orsi et al., 2010; Tasnim and Kelleher, 2018). Despite these fitness costs, nearly all genomes are populated by TEs (reviewed in Huang et al., 2012), and TEs frequently colonize new genomes through horizontal transfer (reviewed in Schaack et al., 2010).

Host regulation of transposition is as ubiquitous as TEs themselves; both prokaryotic and eukaryotic genomes control TE activity through small-RNA-mediated silencing pathways (reviewed in Blumenstiel, 2011). In eukaryotic germlines, small-RNA-mediated silencing of TEs is enacted by complexes of Argonaute proteins and small-interfering RNAs (siRNAs) or Piwi-interacting RNAs (piRNAs). The siRNAs and piRNAs are produced from anti-sense TE transcripts, and identify TE-derived mRNAs by base complementarity. Argonaute proteins then silence target TEs transcriptionally or post-transcriptionally. Consistent with prevention of germline DNA damage, small-RNA-mediated silencing suppresses TE-associated dysgenic sterility in *Drosophila* (Blumenstiel and Hartl, 2005; Brennecke et al., 2008; Chambeyron et al., 2008; Rozhkov et al., 2010) and *Arabidopsis* (Josefsson et al., 2006). Transcriptional silencing may further reduce the rate of ectopic recombination between TE insertions by inducing heterochromatin formation at TE loci (Haynes et al., 2006; Sentmanat and Elgin, 2012; Sienski et al., 2012).

Although the mechanism of small-RNA-mediated silencing has been investigated extensively, we know very little about how it evolves when a genome is invaded by a new TE. In part, this reflects an absence of opportunity, since, although ancient TE invasions are easily identified from comparative genomic data (reviewed in Schaack et al., 2010), ongoing invasions have only rarely been detected in extant populations (Evgen’ev et al., 2000; Hill et al., 2016; Kidwell, 1983; Periquet et al., 1989). Additionally, the adaptive evolution of transpositional repression is a long-standing theoretical challenge in sexually-reproducing organisms. Prior to the discovery of small-RNA-mediated silencing, a series of classic analytical models suggested that positive selection on *trans*-acting transpositional repressors, such as small-RNA-encoding loci, was likely to be weak, because recombination would rapidly decouple repressor alleles from the chromosomal regions that they protect from TE-induced mutation load (Charlesworth and Langley, 1986). Restrictive conditions for the adaptive evolution of repressor alleles are difficult to reconcile with the prevalence of small-RNA-mediated TE silencing across the tree of life.

When a genome is invaded by a new TE, small-RNA-mediated silencing is thought to evolve by transposition of the invading TE into a small-RNA-producing site, thereby initiating the production of new RNA species that target the invading element for silencing (Brennecke et al., 2008; Khurana et al., 2011). TE insertions into small-RNA-producing sites therefore represent a class of repressor alleles that arise via *de novo* mutation during an invasion event. Using forward simulations of retrotransposon invasion into a *Drosophila melanogaster*-like genome, Lu and Clark (2010) demonstrated that small-RNA-producing insertions are targets of positive selection and made a general case for adaptive evolution of small-RNA-mediated TE silencing. However, because their model is not based on any particular TE family, it is unclear how accurately the results reflect an actual invasion event. Additionally, Lu and Clark (2010) did not explicitly consider dysgenic sterility or ectopic recombination, two major TE-imposed fitness costs that could enhance positive selection on repressor alleles.

Here we draw on an extensive body of literature on *P*-element DNA transposons and their recent invasion into natural populations of *D. melanogaster* (reviewed in Kelleher, 2016), to motivate a forward, individual-based simulation model of the evolution of small-RNA-mediated repression in response to a TE invasion event. By incorporating empirically estimated fitness costs, transposition rates, and effects of small-RNA-mediated silencing, our model recapitulates the key features of the *P*-element invasion, including both the spread of the TE through natural populations and the acquisition of repression by the host. We then interrogate the degree to which different TE-imposed fitness costs (*i.e.*, deleterious mutations, dysgenic sterility, and ectopic recombination) enhance positive selection on repressor alleles. Finally, we consider how the distinct genetic architecture of small-RNA-mediated TE silencing—in which many sites across the genome produce small RNAs when occupied by a TE—affects the evolution of repression.

We demonstrate that small-RNA-mediated silencing of transposition alone evolves rapidly and adaptively in response to genome invasion. We further demonstrate that dysgenic sterility and ectopic recombination can greatly enhance the strength of positive selection on insertions in small-RNA-producing sites. Finally, we reveal that the abundance of small-RNA-producing sites in the host genome is a critical parameter that influences both the proliferation of the TE and the adaptive behavior of repressor alleles. Our study provides fundamental insights into selective forces that influence the evolution of repression, as well as an updated framework for modeling TE invasion.

## Model

We employ an individual-based Wright-Fisher model of evolution, with a selection-reproduction-transposition life cycle, constant population size, and discrete, non-overlapping generations. See Table 1 for a list of parameters.

**Table 1.**
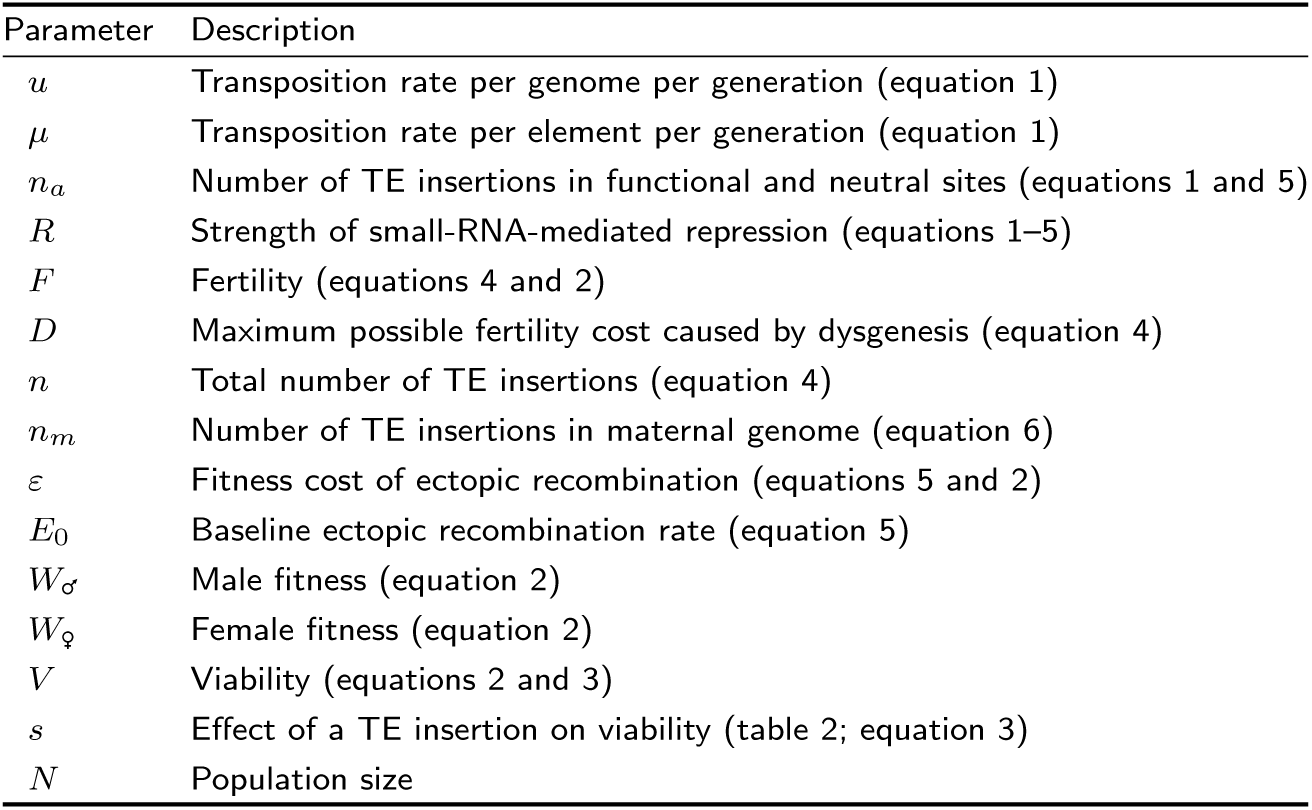
Model parameters

### Genome

The genome is modeled after that of *D. melanogaster*. Each individual is diploid with a haploid complement of 3 chromosomes: one sex chromosome containing 226 sites, and two autosomes containing 442 and 527 sites, respectively (total: 1,195 sites). The numbers of sites per chromosome are based on the lengths of *D. melanogaster* chromosomes X, 2 and 3, with one site every *∼*10^5^ base pairs (we ignore chromosome 4, which accounts for less than 1% of the genome). Likewise, the recombination rates at each chromosomal site are set to reflect the actual *D. melanogaster* recombination rates (Comeron et al., 2012).

Previous theoretical models of TE dynamics assume that TE insertions in all genomic sites are equally deleterious, and that they exhibit negative epistasis with respect to fitness (*e.g.*, Dolgin and Charlesworth, 2008; Lee and Langley, 2012; Lu and Clark, 2010). The latter assumption appears to be required for TE copy number to evolve to equilibrium in the absence of repression (Charlesworth and Charlesworth, 1983). Although it has been proposed that the aneuploid progeny that result from ectopic exchange (Langley et al., 1988; Petrov et al., 2003, 2011), or copy-number dependent heterochromatin formation (Lee, 2015; Lee and Karpen, 2017; Lee and Langley, 2010), could produce negative epistasis between TE insertions, the commonly used mathematical expression for negative epistasis (Charlesworth, 1990) is based on the observed pattern of fitness decline in a mutation accumulation experiment on *D. melanogaster* (Mukai, 1969). The extent to which the results of the Mukai (1969) experiment reflect the fitness consequences of interactions between TE insertions is unknown.

Here, we take a different approach, by considering a more complex and realistic genome in which the fitness effects of TE insertions vary between sites. We furthermore drew extensively on empirical data to determine composition of the genome and the distribution of fitness effects for insertions into individual sites. The genome contains four kinds of sites: small-RNA sites, pseudo-small-RNA sites, functional sites and neutral sites, which differ in their effects on viability, their participation in transposition and ectopic recombination, and their role in small-RNA-mediated silencing. The properties of each type of site are described below.

### Functional sites

Functional sites cause a multiplicative change in viability (equation 3)—and, therefore, fitness equation 2)—when occupied by a TE. Thus, there is no epistasis for viability. We furthermore assume that the effects on viability of TE insertions in functional sites are completely recessive, as has been observed empirically for *P*-elements (Mackay et al., 1992). To determine the number of functional sites in our theoretical genome, we relied on results from extensive *P*-element mediated mutagenesis screens, which indicate that ≳ 80% of new *P*-element insertions occur in gene bodies, within 500 bp of a transcription start site (Bellen et al., 2011, 2004). Accordingly, 80% (956) of sites in our theoretical genome are functional (Table 2) and affect viability when homozygous for a TE insertion.

**Table 2.**
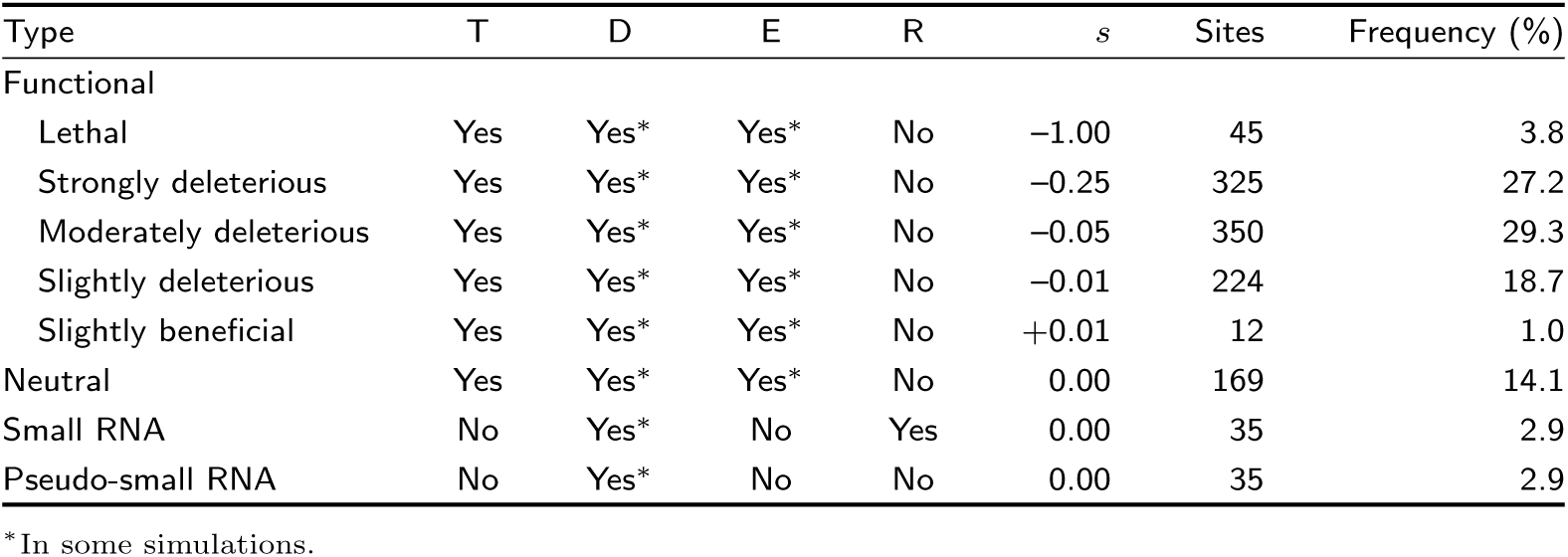
Viability selection coefficients (*s*), and contribution to transposition (T), dysgenic sterility (D), ectopic recombination (E), and repression of transposition (R) of TE insertions in the 1,195 sites of different kinds

| Type | T | D | E | R | $s$ | Sites | Frequency (%) |
| --- | --- | --- | --- | --- | --- | --- | --- |
| Functional |  |  |  |  |  |  |  |
| Lethal | Yes | Yes* | Yes* | No | −1.00 | 45 | 3.8 |
| Strongly deleterious | Yes | Yes* | Yes* | No | −0.25 | 325 | 27.2 |
| Moderately deleterious | Yes | Yes* | Yes* | No | −0.05 | 350 | 29.3 |
| Slightly deleterious | Yes | Yes* | Yes* | No | −0.01 | 224 | 18.7 |
| Slightly beneficial | Yes | Yes* | Yes* | No | +0.01 | 12 | 1.0 |
| Neutral | Yes | Yes* | Yes* | No | 0.00 | 169 | 14.1 |
| Small RNA | No | Yes* | No | Yes | 0.00 | 35 | 2.9 |
| Pseudo-small RNA | No | Yes* | No | No | 0.00 | 35 | 2.9 |

\*In some simulations.

### Neutral sites

169 neutral sites (14.1% of sites in the genome) have no effect on viablity when occupied by a TE. However, TE insertions in neutral sites, like those in functional sites, contribute to both dysgenic sterility and the production of inviable gametes through ectopic recombination in some simulations (see ‘Dysgenic sterility’ and ‘Ectopic recombination’ below).

### Small-RNA sites

35 sites (2.9% of sites in the genome) establish small RNA silencing when occupied by a TE, and represent the piRNA producing regions in the *D. melanogaster* genome known as piRNA clusters (Brennecke et al., 2007). The locations of small-RNA sites are randomly drawn from the known locations of piRNA clusters in the *D. melanogaster* genome (Brennecke et al., 2007). Because piRNA clusters are generally found in heterochromatic regions where chromatin compaction restricts transposition and recombination (Brennecke et al., 2007), TE insertions in small-RNA sites do not transpose and do not participate in ectopic recombination in our model. However, TE insertions in small-RNA sites do contribute to dysgenic sterility in some simulations (see ‘Dysgenic sterility’, below).

### Pseudo-small-RNA sites

35 sites (2.9% of sites in the genome) do not contribute to small RNA silencing but are otherwise indistinguishable from small-RNA sites. Biologically, they represent transcriptionally quiescent heterochromatic sites where ectopic recombination is unlikely. They also provide a neutral site class against which to evaluate the frequency spectra of small-RNA sites. Pseudo-small-RNA sites are placed near telomeres or centromeres, which mirrors the distribution of piRNA clusters in the *D. melanogaster* genome. Each chromosome arm contains three and four pseudo-small-RNA sites at the proximal and distal ends, respectively, skipping sites that are already assigned as either small RNA or functional.

### Transposition, Excision, and Inactivation

The transposition rate per genome per generation is given by

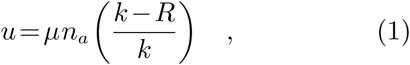

where *µ* is the transposition rate per element per generation of unrepressed TEs, *n*_*a*_ is the number of TEs in the genome that can be mobilized in the presence of transposase (*i.e.*, those in functional and neutral sites, but not in small-RNA or pseudo-small-RNA sites), *R* is the strength of small-RNA-mediated repression (see equation 6), and *k* is a constant. We consider *µ* = 0.1 and *µ* = 0.01 according to the range of empirical estimates for unregulated transposition of *P*-elements (Berg and Spradling, 1991; Eggleston et al., 1988; Kimura and Kidwell, 1994; Robertson et al., 1988). We also set *k* = 1.01, in order to limit the reduction in transposition rate caused by small-RNA-mediated silencing to two orders of magnitude, as is suggested by observed differences in *P*-element activity in the presence and absence of regulation (Eggleston et al., 1988).

Precise *P*-element excisions occur during transposition, with an estimated frequency of 0.15% (per element per genome transposition event) when the insertion is homozygous and *∼*13.5% when the insertion is heterozygous (Engels et al., 1990). We therefore assume that the excision rate is 10% of the transposition rate. Mutational loss of transposase-encoding capacity, which occurs predominantly through internal deletions that are acquired during transposition (Staveley et al., 1995), occurs at a rate of 0.01*µ* per element per generation.

### Fitness

Male and female fitness are given by

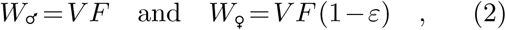

respectively, where

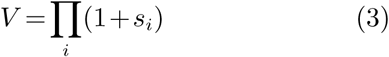

is viability; *s*_*i*_ is the viability selection coefficient of the TE insertion into the *i* locus in a hemizygous or homozygous state; *F* is fertility (equation 4); and *ε* is the fitness cost of ectopic recombination (equation 5).

### Viability

When homozygous, the effect on viability of a TE insertion at a functional site can be lethal (selection coefficient, *s* = *−*1), strongly deleterious (*s* = *−*0.25), moderately deleterious (*s* = *−*0.05), slightly deleterious (*s* = *−*0.01), or slightly beneficial (*s* =+0.01). The frequency of each functional site class in the theoretical genome is described in Table 2 and is based on the empirically estimated distribution of fitness effects of individual *P*-element insertions (Cooley et al., 1988; Lyman et al., 1996; Mackay et al., 1992). Specifically, *P*-element mutagenesis revealed that the frequency of recessive lethal mutations produced by individual insertions is in the range 3.8–10% (Cooley et al., 1988; Mackay et al., 1992). Therefore, we conservatively assigned 3.8% of sites in the theoretical genome to behave as recessive lethals when occupied by a TE. Similarly, the average homozygous viability fitness effect of a TE insertion in the theoretical genome, 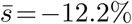, matches a quantitative estimate from a series of *P*-element insertion lines (Mackay et al., 1992).

### Dysgenic Sterility

When *Drosophila* males bearing a transpositionally active TE family are crossed to females who do not produce a sufficient abundance of piRNAs that regulate that TE family, their F1 offspring can suffer from dysgenic sterility (Bingham et al., 1982; Blackman et al., 1987; Brennecke et al., 2008; Bucheton et al., 1984; Evgen’ev et al., 1997; Hill et al., 2016; Yannopoulos et al., 1987). We therefore modeled dysgenic sterility as a function of the number of genomic TEs, whose unregulated activity is thought to induce the sterility phenotype (equation 4). Indeed, in the case of *P*-element induced dysgenic sterility, the proportion of offspring that are sterile has been correlated with the *P*-element germline excision rate, consistent with *P*-element activity as the cause of sterility (Kocur et al., 1986). Dysgenic sterility is further modified in our model by the production of small RNAs in the maternal genotype (*R*).

Dysgenic sterility occurs in both sexes, although it is most extensively studied in females, in whom it tends to be more severe (Kidwell et al., 1977). In particular, Bingham et al. (1982) described the quantitative relationship between the number of paternally inherited *P*-elements and the incidence of dysgenic sterility among F1 female offspring. The paternal genotypes that induced dysgenesis in these crosses were otherwise isogenic, such that differences in dysgenic sterility among F1 females could be exclusively attributed to *P*-element copy number. They observed that *n* = 4 *P*-element copies produced *∼*50% dysgenic offspring, whereas *n* ≳ 9 copies produced ≳ 80% dysgenic offspring. There was very limited data on induction by paternal strains containing *n<* 4 *P*-element copies.

We matched these empirical observations by expressing the fertility, *F*, of an individual as a function of its total diploid TE copy number (*n*). In contrast to viability fitness therefore, where the fitness effects of individual insertions are recessive within loci, and additive across loci, each *P*-element insertion contributes epistatically to the fitness cost of ectopic recombination, regardless of whether it is found in a homozygous or heterozygous state. Fertility is given by

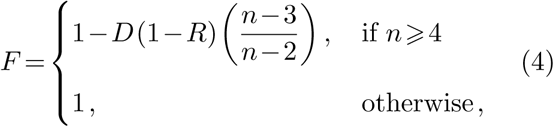

where *D* is a constant that represents the maximum fertility cost, and can be varied from 0 (no dysgenic sterility) to 1 (complete dysgenic sterility) and *R* is the strength of small-RNA-mediated repression in the maternal genotype (see equation 6). *F* can vary from 0 (complete infertility) to 1 (complete fertility).

### Ectopic Recombination

Ectopic recombination occurs between TE insertions in different genomic sites, and produces inviable gametes in our model. We assume that the fitness cost of ectopic recombination is proportional to the number of pairwise combinations of TEs in the genome, excluding insertions in small-RNA and pseudo-small-RNA sites. The fitness consequences of ectopic recombination are, therefore, analogous to the negative epistasis between TE insertions that is assumed by previous models (Dolgin and Charlesworth, 2006, 2008; Lee and Langley, 2012; Lu and Clark, 2010). Each *P*-element insertion contributes epistatically to the fitness cost of ectopic recombination, regardless of whether it is found in a homozygous or heterozygous state.

Small-RNA-mediated silencing acts as a multiplicative modifier of ectopic recombination, which reflects the potential suppression of recombination at TE loci due to heterochromatin formation (Haynes et al., 2006; Klenov et al., 2007). The fitness cost of ectopic recombination (*ε*) is described by the following expression:

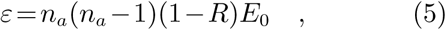

where *E*_0_ is the baseline ectopic recombination rate; *n*_*a*_ is the number of TE insertions that can contribute to ectopic recombination (*i.e.*, do not occur in small-RNA or pseudo-small-RNA sites); and *R* is the strength of small-RNA-mediated repression.

### Small-RNA-Mediated Silencing

piRNA-mediated silencing represses both germline TE activity and dysgenic sterility in *Drosophila* (Brennecke et al., 2008; Khurana et al., 2011). The production of regulatory piRNAs requires the presence of at least one homologous TE insertion in a piRNA cluster, and is amplified by the presence of additional TE copies elsewhere in the genome (Brennecke et al., 2008; Zanni et al., 2013). The enhancement of piRNA production by TE insertions outside of piRNA clusters is explained by the feed-forward biogenesis of piRNAs through transcriptional silencing of sense TE transcripts (*i.e.* ping-pong biogenesis, Brennecke et al. 2007; Gunawardane et al. 2007). piRNAs are maternally transmitted, such that piRNA production in the mother determines the strength of silencing in the offspring (Brennecke et al., 2008).

Empirical data regarding the quantitative relationship between TE copy numbers inside and outside of piRNA clusters and reductions in transposition rate are sparse for *P*-elements, and entirely lacking for other TE families. We therefore harnessed the extensive data on piRNA-mediated repression of dysgenic sterility (Marin et al., 2000; Ronsseray et al., 1991, 1996, 1998; Simmons et al., 2014, 2007, 2012; Stuart et al., 2002) (Supplemental Material Table S1, figure S1). Based on these observations, we describe the strength of small-RNA-mediated silencing, *R*, conferred by maternal genotypes with at least one occupied small-RNA site as

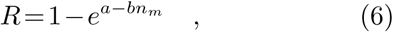

where *n*_*m*_ is the total number of TE copies in the maternal genome, and *a* and *b* are constants. We fit equation 6 by nonlinear least squares to the data in Table S1 and obtained: *a* = 0.8*±*0.5 and *b* = 1.0*±*0.4 (estimates and 95% confidence intervals). *R* can vary from 0 (no repression) to 1 (complete repression).

### Evolution

Each population is defined by the following parameters: population size (*N*), transposition rate (*µ*), baseline ectopic recombination rate (*E*_0_), and the maximum possible fertility cost caused by dysgenic sterility (*D*). In the initial population, the sex of each individual is decided randomly, so that the expected sex ratio is 1:1. The range of parameter values we considered over our simulations is described in Table 3. 5% of the population harbors one TE insertion at a randomly chosen neutral site on chromosome 2. The remaining 95% of the population harbors no TE insertions.

**Table 3.** Range of parameter values explored for diploid population size (*N*), unrepressed transposition rate (*µ*), maximum possible fertility cost caused by dysgenic sterility (*D*), baseline ectopic recombination rate (*E*_0_)

| Parameter | Symbol | Values | Units |
| --- | --- | --- | --- |
| diploid population size | $N$ | $2 \times 10^2, 2 \times 10^3, 2 \times 10^4$ | individuals |
| transposition rate | $\mu$ | $10^{-1}, 10^{-2}$ | insertions per element per generation |
| maximum dysgenic sterility | $D$ | 0, 0.1, 0.25, 0.5, 1 | proportional fitness |
| ectopic recombination rate | $E$ | 0, $10^{-1}, 10^{-2}, 10^{-3}, 10^{-4}, 10^{-5}$ | proportional fitness |

We simulated reproduction by randomly selecting one male and one female, each with probability proportional to their fitness (equation 2). After an individual parent is selected to reproduce, the number of new TE copies to transpose within its germline is drawn from a Poisson distribution with parameter *u* (equation 1). The new TE copies are distributed randomly among unoccupied sites and are only used to produce one specific offspring; if a parent produces multiple offspring, transposition is repeated independently for each offspring produced.

Haploid gametes are produced by meiosis, simulated by choosing one chromosome per pair at random. Crossing over occurs in females only, as is true for *D. melanogaster* (Morgan, 1914). We assumed no crossover interference. Diploid progeny are formed by fusion of one gamete from each parent. Once *N* offspring are produced, the parents are discarded.

## Results

### Simulated invasions recapitulate the historical record

We began by considering a conservative model, in which TEs impose fitness costs only by occupying functional sites in the host genome, and repression benefits the host by preventing the occurrence of these deleterious insertions. Dysgenic sterility and ectopic recombination were not incorporated in the model (*D* = 0*, E*_0_ = 0). We compared two baseline transposition rates *µ* = 0.1 and *µ* = 0.01, which encompass the range of most empirically estimated values for unregulated *P*-element transposition (Berg and Spradling, 1991; Bingham et al., 1982; Eggleston et al., 1988; Kimura and Kidwell, 1994; Robertson et al., 1988).

Simulations were started from populations of *N* = 2,000 in which 5% of individuals harbor a single autosomal TE insertion at a neutral site. Populations were then allowed to evolve for 2,000 generations. At the higher transposition rate of *µ* = 0.1, the TE successfully invaded in 100% of simulations (out of 100 runs; figure 1A, red). This occurred very rapidly, with 100% of genomes harboring at least one TE copy within 100 generations. By contrast, at the lower transposition rate of *µ* = 0.01 only 71% of simulations exhibited successful invasions within 2,000 generations (out of 100 runs; figure 1A, black). The remaining 29% of simulations lost the *P*-element entirely within 2,000 generations, presumably due to drift and negative selection. The two transposition rates further differ in the average copy number exhibited by individual genomes throughout the invasion process (figure 1B), with copy numbers stabilizing around 50 copies/genome under the higher transposition rate, but only approximately 30 copies/genome under the lower transposition rate.

**Fig. 1.**
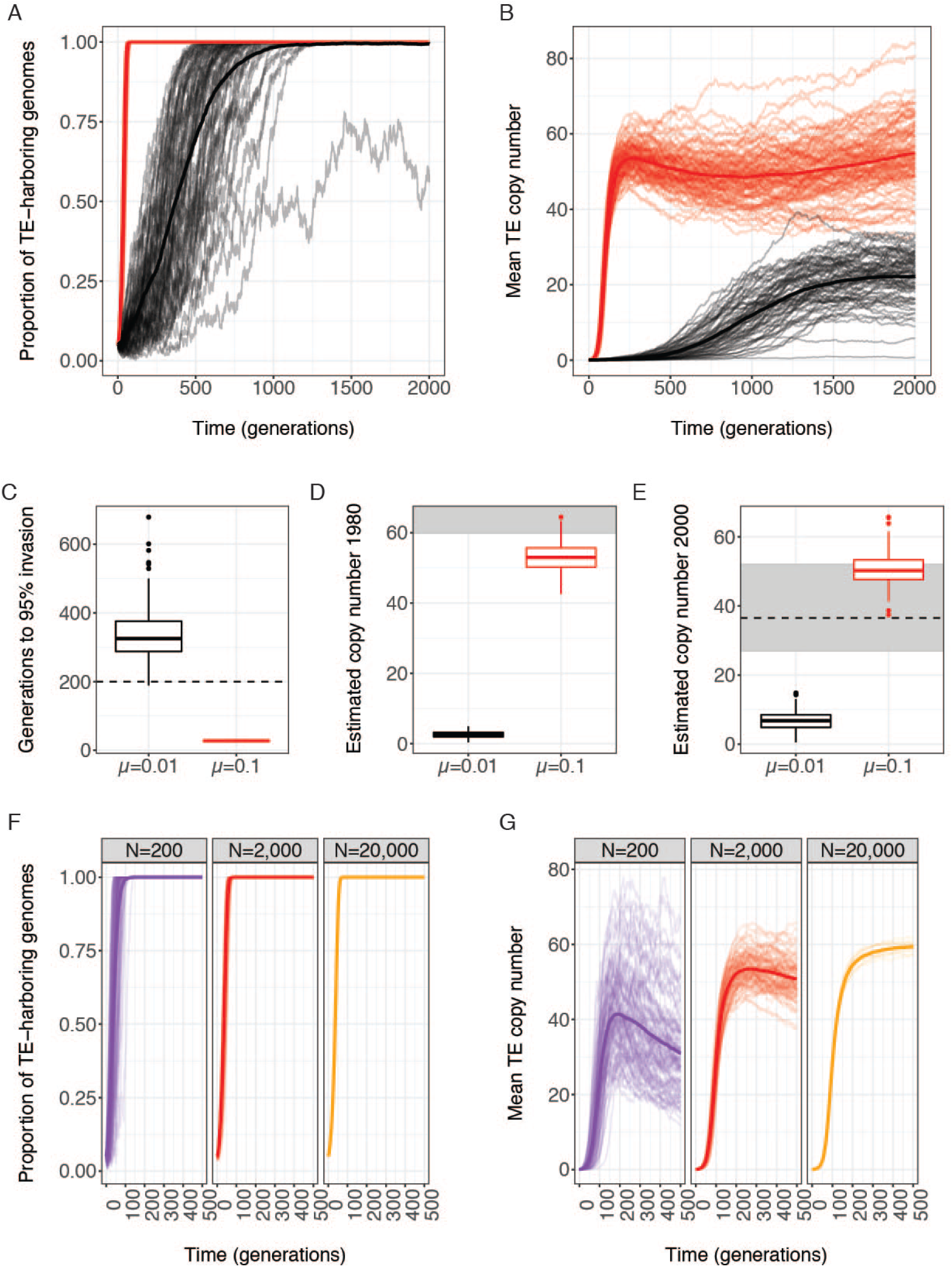
Rapid population invasion requires a high transposition rate. Evolution of the A) proportion of individuals with at least one genomic TE and B) mean diploid TE copy number (*n*) over 2,000 generations in 100 replicate simulated populations of *N* = 2,000 individuals are shown. C) The number of generations required for the proportion of individuals with at least one genomic TE to increase from 33% to 95%. D) and E) The mean diploid copy number in simulated populations after 300 generations D) and 500 generations E) after 1950 (*i.e.* first generation where *>* 33% of individual genomes carry TEs). Gray boxes indicate empirical estimates of historic values (Bingham et al., 1982; Ronsseray et al., 1989; Srivastav and Kelleher, 2017). For panels A-E, the trajectory of invasion is compared for two different baseline transposition rates: *µ* = 0.1 (red) and *µ* = 0.01 (black). F) proportion of individuals with at least one genomic TE and G) mean diploid TE copy number (*n*) over 500 generations in 100 replicate simulated populations of *N* = 200 and *N* = 2,000 individuals, and 10 replicate simulated populations of *N* = 20,000 individuals are compared. Individuals in all simulations (A–G) experienced neither dysgenic sterility nor ectopic recombination (*D* = 0*, E*_0_ = 0).

In light of the difference in invasion dynamics between the two transposition rates, we wondered which set of simulations more closely matched the historical record of the *P*-element invasion. We focused on American populations (North and South), for which the historical record of the *P*-element invasion is particularly comprehensive (reviewed in Kelleher, 2016). With the exception of a single strain collected in 1938, all wild-derived strains collected prior to 1950 lack genomic *P*-elements, indicating the *P*-element was quite rare (Anxolabéhère et al., 1988; Bingham et al., 1982). By contrast, *∼*33% of collections from the 1950s (4/12 strains) and 1960s (4/13 strains), contain *P*-elements, and *∼* 100% of *>* 200 strains collected after 1971 contain *P*-elements (Anxolabèhére et al., 1988). Thus, in *<* 20 years, and possibly *<* 10, *P*-elements went from intermediate frequency to complete fixation in North America, a rapid colonization event that was recapitulated in other geographic locations (Anxolabéhère et al., 1988).

To compare the observed colonization rate to our simulated data, we assumed that the first simulated generation where *>* 33% of genomes contained *P*-elements corresponded to the year 1950. We then determined how many additional generations were required for *>* 95% of genomes to contain *P*-elements. Assuming 10 generations per year for 20 years, 95% colonized genomes should be achieved in *<* 200 generations. Indeed, this occurred in 100% of simulations where *µ* = 0.1, but only 7% of simulations where *µ* = 0.01 (both out of 100 runs; figure 1C). Thus, the speed of the invasion in natural population seems more consistent with *µ* = 0.1 than *µ* = 0.01.

We also considered the historical record with regards to *P*-element copy numbers in North American genomes. *P*-element containing strains collected in the late 1960s through the early 1980s were estimated by two different studies to contain 60–100 copies/diploid genome, based on Southern blotting (Bingham et al., 1982) and *in situ* hybridization of polytene chromosomes (Ronsseray et al., 1989). Based on next generation sequencing data, Srivastav and Kelleher (2017) estimated somewhat lower values of *∼*38–52 copies per diploid genome from a panel of isofemale lines collected in North Carolina in 2004 (Srivastav and Kelleher, 2017). It should be noted that these values do not necessarily suggest a decrease in *P*-element copy number since the 1980s, because estimates from Southern blotting may be somewhat inflated by polymorphism, while estimates from deep-sequencing data are likely to be somewhat low, due to unannotated heterochromatic insertions (Zhang and Kelleher, 2017), challenges in estimating copy number across structural variants (Srivastav and Kelleher, 2017), and loss of deleterious copies during inbreeding.

We compared our simulated copy numbers to historical values from 1980 and 2004 by determining the mean diploid copy number in each simulated data set 300 and 500 generations after 33% of individual genomes contain *P*-elements (year 1950). For the higher transposition rate, the mean estimated copy numbers for 1980 and 2000 were 53 and 50 in the simulated data, with many or most simulation data sets falling within the range of historical estimates (figure 1D,E red). By contrast, the estimated copy numbers at lower transposition rates are an order of magnitude too low, with mean estimated copy number of 2.5 for 1980 and 6.6 for 2000, and 0% of simulated data sets (out of 100) exhibiting copy numbers that are consistent with historical estimates (figure 1D,E black).

Collectively, our observations suggest that invasion models with the higher transposition rate of *µ* = 0.1 closely recapitulate the historical invasion of *P*-elements in natural populations, in terms of the rate of spread of the element and the genomic copy numbers observed. By contrast, simulations with lower transposition rates of *µ* = 0.01 exhibit unrealistically slow invasions with with copy numbers that are more than an order of magnitude lower than historical values. Thus, the transposition rate appears to be a key parameter in the invasion behavior of the element in our model. This observation is robust to different population sizes, as simulations of larger (*N* = 20,000) and smaller (*N* = 200) populations exhibit similar behavior in terms of invasion time (figure 1F). Furthermore, although mean copy numbers are slightly higher in larger populations (figure 1F), the differences between *N* = 2,000 and *N* = 20,000 are so modest that realistic population sizes for *D. melanogaster* (*e.g.*, *N_e_ ≈* 2*×*10^6^, Garud et al., 2015) are not expected to differ dramatically in this respect.

### Deleterious viability fitness effects are sufficient to select for host repression

Although TE copy numbers increase faster than linearly for the first *∼* 100 generations in simulations where *µ* = 0.1 (figure 1B, 2A, red), this growth slows over the subsequent *∼* 100 generations, stabilizing around generation 200 at *∼* 52 copies per genome. Reduced TE proliferation is most likely explained by the evolution of maternal repression. Mean maternal repression (*R*) increased rapidly in simulated populations, and approached complete repression after only 200 generations (figure 2A, blue). By contrast, when repression was not allowed to evolve (*i.e.*, *R* = 0 for all individuals, regardless of genotype), 100% of simulated populations went extinct within 173 generations (out of 802 runs) because all individuals of one sex were homozygous or hemizygous for a lethal TE insertion. For 795 of the runs, no viable males remained, suggesting hemizygous X-chromosomes carrying recessive lethals contributed disproportionately to population extinction.

**Fig. 2.**
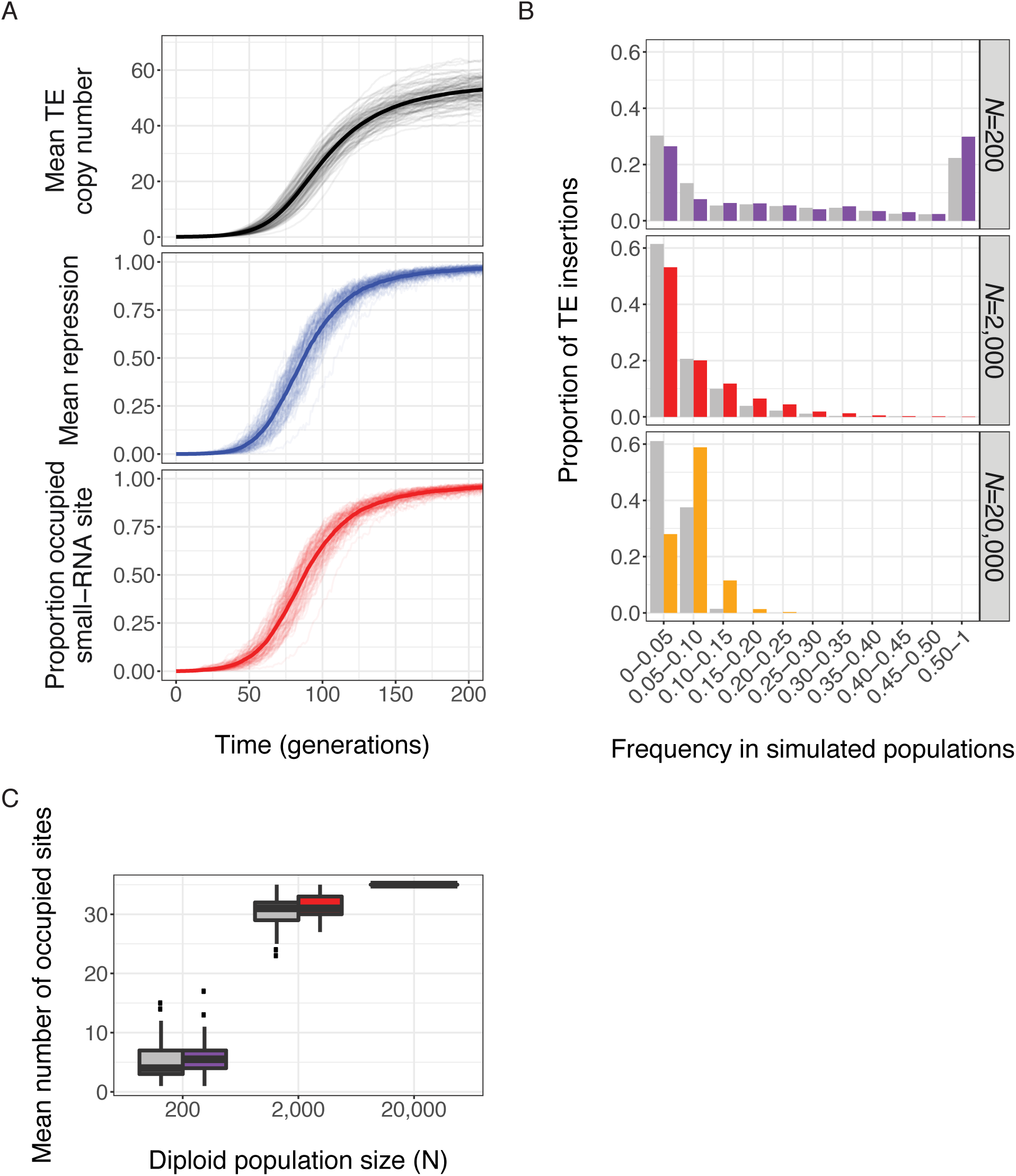
Invasion is followed by rapid and adaptive evolution of host repression. A) Evolution of mean genomic TE copy number (red, top), the mean maternal repression (*R*, blue, middle) and proportion of the population with *≥* 1 occupied small-RNA site (black, bottom) over 200 generations in 100 replicate simulated populations of *N* = 2,000 individuals with a baseline transposition rate *µ* = 0.1. Thick lines represent the average responses across all replicate populations. The site frequency spectra of occupied small-RNA-producing sites (red), which enact maternal repression of transposition, and pseudo-small-RNA-producing sites (gray) which do not repress transposition. B) Results are shown for after 500 generations for simulations with a baseline transposition rate of *µ* = 0.1. Individuals in all simulations experienced neither dysgenic sterility nor ectopic recombination (*D* = 0*, E*_0_ = 0).

In our model, repression of transposition requires that at least one small-RNA-producing site in the maternal genome is occupied by a *P*-element insertion. Therefore, the spread of the maternal repressive phenotype (*R*) is accompanied by an increasing frequency of individuals with *P*-element insertions in small-RNA-producing sites (figure 2A, black). To examine the role of positive selection in the spread of these repressor alleles, and more broadly in the evolution of repression itself, we considered the site frequency spectrum of TE insertions in small-RNA producing sites after 500 generations of simulated evolution. We compared these sites to control pseudo-small-RNA sites, neutral sites that do not produce small RNAs when occupied by TEs but otherwise are matched to small-RNA sites in their genomic locations and fitness effects (see Model). Positive selection will increase the frequency of derived alleles in the population as compared to a neutral model (Fay and Wu, 2000; Nielsen, 2005).

Consistent with positive selection for repression, the site-frequency spectrum of occupied small-RNA-producing sites is significantly different than that of pseudo-small-RNA sites (Kolmogorov-Smirnov test, *N* = 200: *D* = 0.10, *p* = 0.01; *N* = 2,000: *D* = 0.09, *p* = 2.14*×*10^*−*15^; *N* = 20,000: *D* = 0.34, *p<* 10^*−*15^), with occupied small-RNA sites segregating at higher frequency than their neutral counterparts across a range of population sizes (figure 2B). Furthermore, the difference in the site-frequency spectra of selected and neutral sites becomes more pronounced in larger populations where selection is more effective. This indicates that positive selection on occupied small RNA sites is not an artifact of modestly enhanced linkage-disequilibrium between small RNA sites and functional sites in smaller populations (Supplemental Material figure S2).

Additional evidence of positive selection is provided by the number of segregating (*i.e.* occupied in at least one individual) small-RNA sites, as compared to pseudo-small-RNA sites, in each simulated population at generation 500. In populations of *N* = 200 and *N* = 2,000, there were a greater numbers of segregating occupied small-RNA sites than pseudo-small-RNA sites (*N* = 200: small RNA 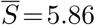, pseudo-small RNA 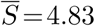, permutation *p* = 0.0067; *N* = 2,000: small RNA 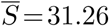, pseudo-small RNA 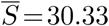, permutation *p* = 0.0036; figure 2C). Because these sites occur in equal numbers in the genome, this indicates that small-RNA-producing insertions are retained by positive selection. The number of segregating small RNA and pseudo-small RNA sites could not be compared for the largest population size (*N* = 20,000), because all sites of both classes were occupied in every replicate population (Figure 2C).

### Moderate dysgenic sterility enhances positive selection for repression

We next expanded our model to consider the effects of dysgenic sterility. In *Drosophila*, dysgenic sterility occurs in the germline of F1 offspring from paternal genotypes containing transpositionally active elements and maternal genotypes that do not produce sufficient regulatory piRNAs (Blumenstiel and Hartl, 2005; Brennecke et al., 2008; Chambeyron et al., 2008; Josefsson et al., 2006; Rozhkov et al., 2010). We therefore modeled fitness costs of dysgenic sterility as dependent on an individual’s TE copy number and the strength of maternal small-RNA silencing. Although our copy-number-dependent function for dysgenic sterility is based on empirical data (Bingham et al., 1982), it is difficult to identify a realistic magnitude for the fertility cost, because the penetrance of the sterility phenotype depends greatly on temperature (Engels and Preston, 1979; Kidwell et al., 1977). We therefore considered a range of values for the maximum reduction in fertility that occurs due to dysgenic sterility (*D* = 0, 0.1, 0.25, 0.5, and 1).

The TE failed to invade completely in 100% (*i.e.* the TE did not occur in *≥* 99% of individual genomes at generation 500) of 377 simulated populations where *D* = 1, and 74% of 381 simulated populations where *D* = 0.5 (Supplemental Material figure S3). Invasions were successful in the remaining 26% of simulations where *D* = 0.5, and in 100% of simulations with lower rates of dysgenic sterility (*D* = 0.1 or 0.25). Failed invasions in the presence of severe dysgenic sterility (*D≥* 0.5) reflect enhanced negative selection against TEs. Similarly, although successful invasions occurred when dysgenic sterility was less costly (*D<* 0.5), they were characterized by reduced TE copy numbers (figure 3A). Importantly, however, the overall timing of these invasions was not altered, as TE copy numbers exhibited the same pattern of early exponential increase followed by stabilization around generation 200 (figure 3A). Therefore, the rapid population invasion that occurs at high transposition rates (*µ* = 0.1, figure 1A,C) is not strongly dependent on the fitness costs of the TE.

**Fig. 3.**
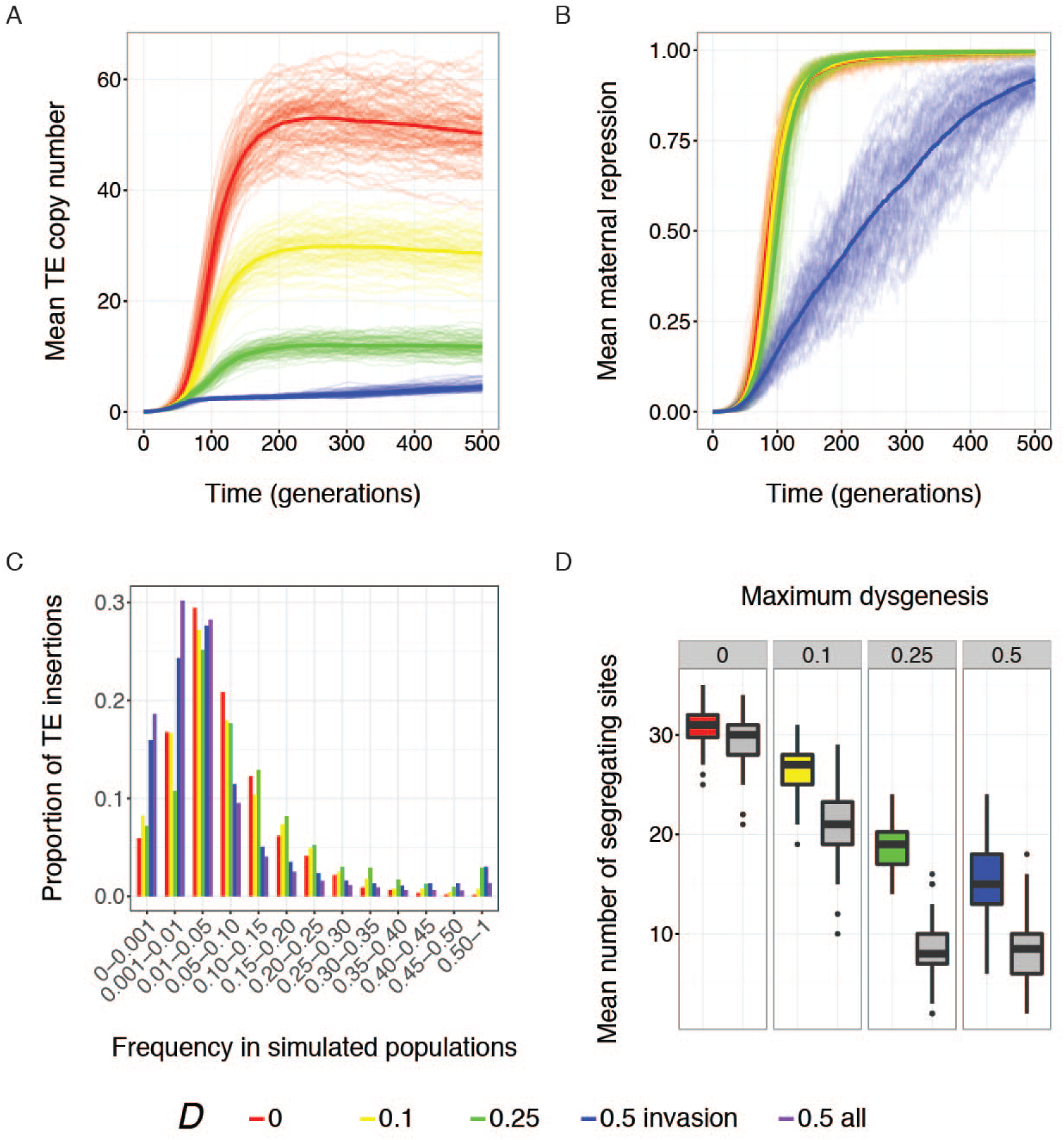
Positive selection on repressor alleles is enhanced by increasing costs of hybrid dysgenesis. Evolution of A) mean diploid TE copy number (*n*) and B) maternal repression (*R*) over 500 generations of simulated evolution. The thick line represents the average response across all replicate populations. C) The site frequency spectra of occupied small-RNA-producing sites after 500 generations of simulated evolution. The spectra were calculated by pooling the data from all replicate simulated populations. Site frequency bins include sites whose frequencies are higher than the lower bound, and less than or equal to the upper bound. D) The average number of occupied small-RNA and pseudo-small-RNA sites in each replicate simulated population after 500 generations. A site is considered occupied if it carries a TE insertion in at least one individual in a population. Population sizes were *N* = 2,000 for all simulations, and individuals experienced no ectopic recombination (*E*_0_ = 0). Colors indicate simulations with different values for maximum hybrid dysgenesis (*D*). For *D* = 0.5, blue indicates only simulated populations that exhibited successful invasion while purple indicates all simulated populations.

Increased negative selection against TEs has potentially opposing effects on the evolution of small-RNA-mediated repression. Although the additional fitness costs imposed by dysgenic sterility could enhance positive selection on occupied small-RNA sites, lower TE copy numbers correspond to a lower rate at which repressor alleles are produced by transposition. With moderate costs of dysgenic sterility (*D<* 0.5), we detect these opposing effects. Although repression evolves rapidly, within *∼*200 generations of invasion, repressive individuals emerge later in the invasion when values of *D* are higher (figure 3B, Supplementary Material figure S3). Nevertheless, higher values of *D* are also associated with more high-frequency occupied small-RNA sites after 500 generations of simulated evolution, indicative of enhanced positive selection for repression (figure 3C). Enhanced positive selection is also evident from the increased number of segregating small-RNA sites, as compared to pseudo-small-RNA sites, in simulated populations (figure 3D).

When the costs of dysgenic sterility were high (*D* = 0.5), we observed a different selective regime, in which TE proliferation was limited by purifying selection to such an extent that positive selection for repression was weakened. In these simulations, the average strength of maternal repression increased in a slower, more linear fashion (figure 3B, blue), and occupied small-RNA sites were not segregating at higher frequency after 500 generations of simulated evolution when compared to *D* = 0.25 (figure 3C, blue versus green). Furthermore, if we consider the full set of simulations for *D* = 0.5 (both successful and failed invasions), high frequency occupied small-RNA sites (*≥* 50% of chromosomes) are even less common, suggesting that the requirement for successful invasion within 500 generations results in an ascertainment bias towards simulated populations with stronger repression (figure 3C, purple).

In addition to a weaker footprint of positive selection on their site frequency spectra, we also find less evidence for selective retention of occupied small-RNA sites in simulated populations when *D* = 0.5. While the mean number of segregating pseudo-small-RNA sites in simulated populations at generation 500 does not differ between *D* = 0.5 and *D* = 0.25 (*t*_198_ = 0.14, *p* = 0.89), larger numbers of segregating small-RNA sites are observed when *D* = 0.25 (mean= 18.67) than when *D* = 0.5 (mean= 15.63, *t*_198_ = 7.09, *p* = 2.35*×*10^*−*11^). This pattern contrasts increased retention of occupied small-RNA sites with increasing costs of dysgenic sterility when *D≤* 0.25 (figure 3D, red, yellow, and green versus gray).

### Ectopic recombination enhances positive selection for repression

We lastly considered the impact of fitness costs imposed by ectopic recombination on invasion success and selection for repression. In our model, the probability of an ectopic recombination event in an individual’s germline increases faster than linearly with TE copy number (equation 5), effectively producing negative epistasis between TE insertions. Thus, this fitness model is most similar to the model employed by previous forward simulations of the evolution of host repression (Lee and Langley, 2012; Lu and Clark, 2010). However, in our simulations small RNA silencing repressed not only transposition, but also the fitness costs associated with ectopic recombination. For simplicity, individuals did not experience hybrid dysgenesis in these simulations (*D* = 0). Similar to other simulations that include small-RNA-mediated silencing (figures 1B, 2A, 3A), we observed an initial exponential increase in TE copy number that tapered off in less than 200 generations (figure 4A). Additionally, higher baseline ectopic recombination rates (*E*_0_), were associated with lower TE copy numbers throughout the simulation (figure 4A), as expected from the higher fitness cost for each TE. Similar to hybrid dysgenesis, fitness costs imposed by ectopic recombination were associated with enhanced positive selection on occupied small-RNA sites. The site frequency spectra for occupied small-RNA sites at the end of the 500 generations of simulated evolution included a greater proportion of high-frequency variants, with 45% of 100 populations exhibiting a high-frequency (*≥* 50%) occupied small-RNA site at the highest baseline ectopic recombination rate we considered (*E*_0_ = 0.01, figure 4B, purple). Similarly, the difference between the average number of occupied small-RNA sites and pseudo-small-RNA sites was more pronounced at higher *E*_0_ values, indicating that selection increasingly retained occupied small-RNA sites (figure 4C).

**Fig. 4.**
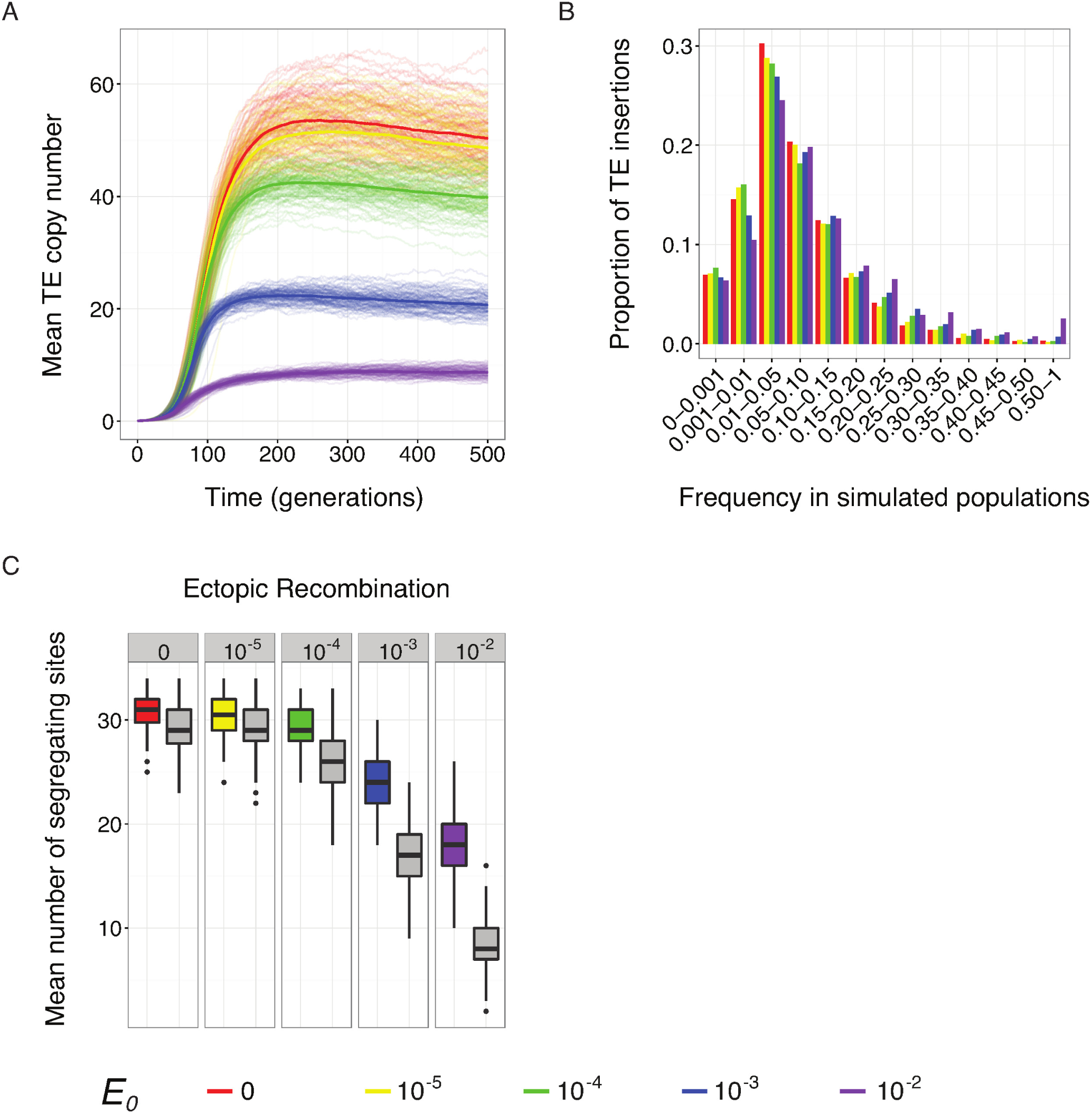
Positive selection on repressor alleles is enhanced by increasing costs of ectopic recombination. Evolution of mean diploid TE copy number (*n*) over 500 generations of simulated evolution. The thick line represents the average response across all replicate simulated populations. B and C) The site frequency spectra of occupied small-RNA-producing sites (B) and the average number of occupied small-RNA and pseudo-small-RNA sites (C) after 500 generations of simulated evolution. Site frequency spectra (B) were calculated by pooling the data from all replicate simulated populations. Site frequency bins include sites whose frequencies are higher than the lower bound, and less than or equal to the upper bound. A site is considered occupied (C) if it carries a TE insertion in at least one individual in a population. Population sizes were *N* = 2,000 for all simulations, and individuals experienced no dysgenic sterility (*D* = 0). Colors compare simulations with different values of the baseline ectopic recombination rate (*E*_0_).

### Large mutational targets reduce positive selection on repressor alleles

Although occupied small-RNA-producing sites were targets of positive selection in many of our simulations, these variants were rarely fixed after 500 generations (figures 2B, 3C, 4B). Rather, a given population was generally segregating for multiple, and sometimes many, occupied small-RNA-producing sites (figures 2C, 3D, 4C). These observations suggest that the evolution of small-RNA-mediated TE silencing is not mutation-limited. Rather, repression may evolve adaptively through a soft-sweep-like process, in which multiple small-RNA-producing TE insertions at different genomic sites increase in frequency by positive selection.

To further interrogate the impact of mutation limitation on the evolution of small-RNA-mediated repression, we considered genomes with different numbers of small-RNA sites. Differences in the size of the mutational target are analogous to different rates of transposition into into small-RNA sites, *i.e.* differences in the individual adaptive mutation rate (*U*_*A*_). Indeed, by estimating *U*_*A*_ as the product of individual transposition rate *U* and the proportion of small sites in the genome we confirmed that *U*_*A*_ is higher throughout the simulation when small sites are common, as compared to when small RNA sites are rare (figure 5A). Furthermore, these differences in the adaptive mutation rate are associated with different trajectories for maternal repression within simulated populations (figure 5B). When small-RNA sites are common, repression is more common early in the invasion (*<* 50 generations), but increases in abundance at a slower exponential rate. In contrast, when there are few small-RNA-producing sites, repression remains rare for much longer, but spreads much more rapidly. Importantly, regardless of the mutational target size, repression evolves rapidly, within 200 generations of TE invasion, indicating the evolution of small-RNA-mediated repression is robust to the abundance of small RNA sites in the genome.

**Fig. 5.**
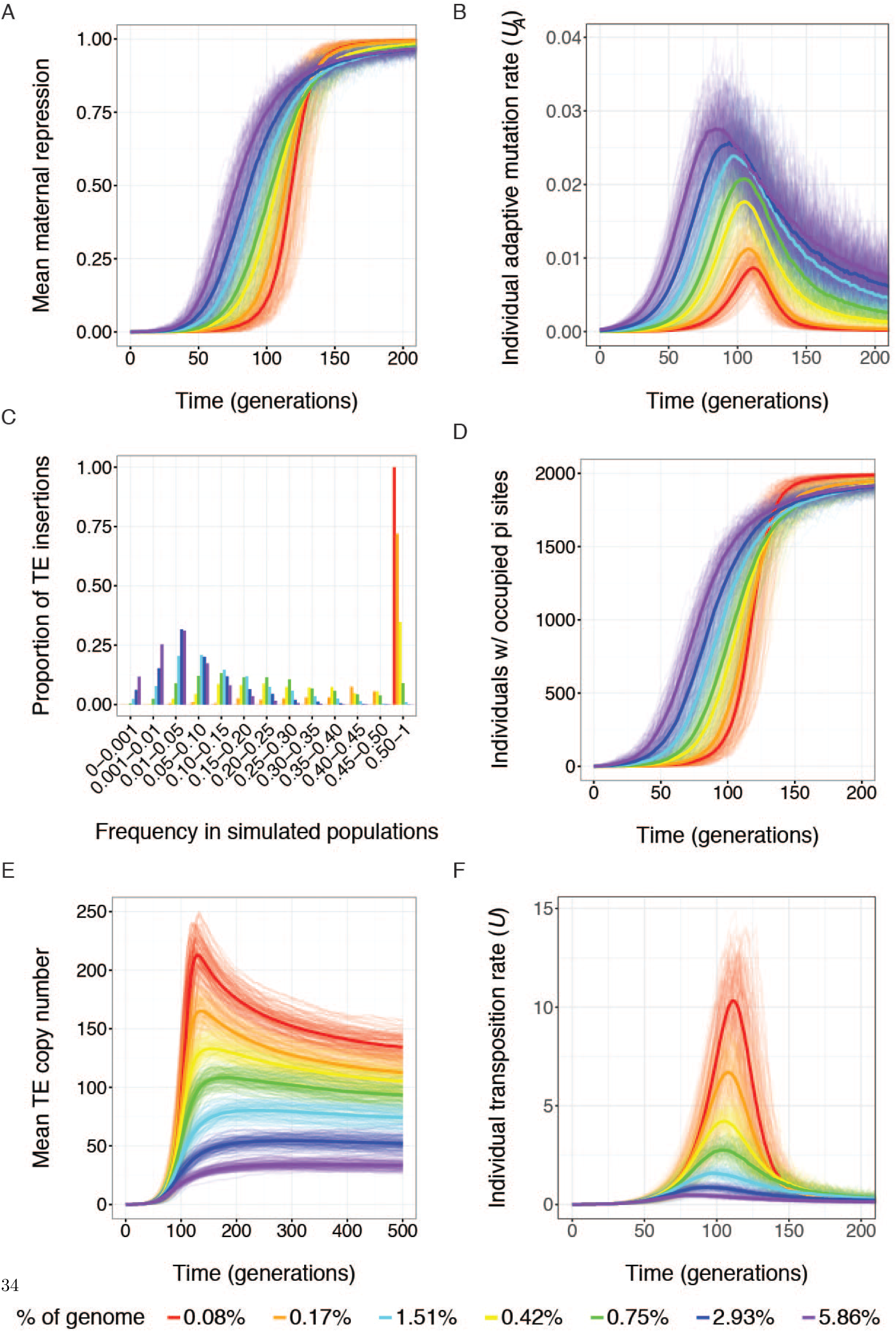
The abundance of small-RNA sites is related to the strength of positive selection for repression. A), B), D) and F) Evolution of the mean individual adaptive mutation rate (*U*_*A*_, insertions into small-RNA sites per genome) (A) the mean maternal repression (*R*) (B) the cumulative frequency of individuals with at least one occupied small-RNA site (D), and the mean individuation transposition rate (*U* insertions per genome)(F) over 200 generations in replicate simulated populations. E) Mean diploid TE copy number (*n*) over 500 generations in simulated populations. The thick line represents the average response across 100 replicate populations. F) The site frequency spectra of occupied small-RNA-producing sites after 500 generations of simulated evolution. Site frequency bins include sites whose frequencies are higher than the lower bound, and less than or equal to the upper bound. The site frequency spectra were calculated by pooling the data from all 100 replicate simulated populations. All simulations included *N* = 2,000 individuals, who experienced neither dysgenic sterility nor ectopic recombination (*D* = 0*, E*_0_ = 0). Colors indicate simulations that involved genomes with different fractions of small-RNA-producing sites.

The more rapid increase in repression when small-RNA sites are rare suggests enhanced positive selection. Although the site frequency spectra of occupied small-RNA sites include more high frequency variants when the mutational target is small (figure 5C), the inference of stronger positive selection is confounded by the fact that occupied small-RNA sites are produced less frequently in more mutation-limited genomes, increasing the probability that the first occupied small-RNA site reaches fixation before a new one arises. We therefore considered the cumulative frequency of individuals harboring at least one insertion in a small-RNA-producing site over the course of simulated evolution (figure 5D). In this way we examine the increase in frequency of the repressor allele class as a whole, independently of the number of alleles. The cumulative frequency of individual genomes with at least one repressor allele increases more rapidly when the mutational target to repression is small, suggesting stronger positive selection.

Stronger selection on repressor alleles in genomes with fewer small-RNA sites is explained by the accumulation of a greater number of TE copies, and by association, an increased individual transposition rate *U*. (figure 5E,F). The benefits of repression are therefore enhanced by an increased deleterious mutation rate, as well as an increased likelihood that deleterious insertions are homozygous and therefore exposed to selection. Indeed, the accumulation of TE copies in genomes with smaller mutational targets is associated with decreased mean population fitness and increased variance in fitness, indicating stronger selection (Supplemental Material figure S5). Although we did not explore whether copy-number equilibrium depends on the size mutational target, we did observe that copy number differences between simulations with genomes containing 0.08% and 5.86% small-RNA sites persisted through 2,000 generations. Simulated populations in which the mutational target was larger (5.86%) exhibited more than *>*3-fold lower diploid copy numbers (36.58) when compared to those with small mutational targets (mean 0.08%= 120.43 *p<* 10^*−*15^).

The inverse relationship between the abundance of small RNA sites and positive selection on maternal repression is also observed at larger population sizes of *N* = 20,000 (Supplementary Material Figure S6). Puzzlingly however, larger populations do not exhibit stronger repression earlier in the invasion process (Figure 6). Under strict mutation-limitation, stronger repression early in the invasion would be predicted in larger populations, because the probability that a repressive individual arises in any given generation increases with population size (Pennings and Hermisson 2006b). However, it must be considered that selection for repression of an invading TE is temporally dynamic, and increases over time as TE copies accumulate. Therefore, early in the invasion process, repressor alleles will be neutral because deleterious insertions are absent or minimal. By the time repressor alleles enjoy a selective advantage, they will be present in similar frequency regardless of population size, but selection will be more effective in larger populations. This mirrors perfectly our observations, in which repression increases at equivalent rates early in the invasion process, regardless of population size, but increases more quickly in later generations in larger populations.

**Fig. 6.**
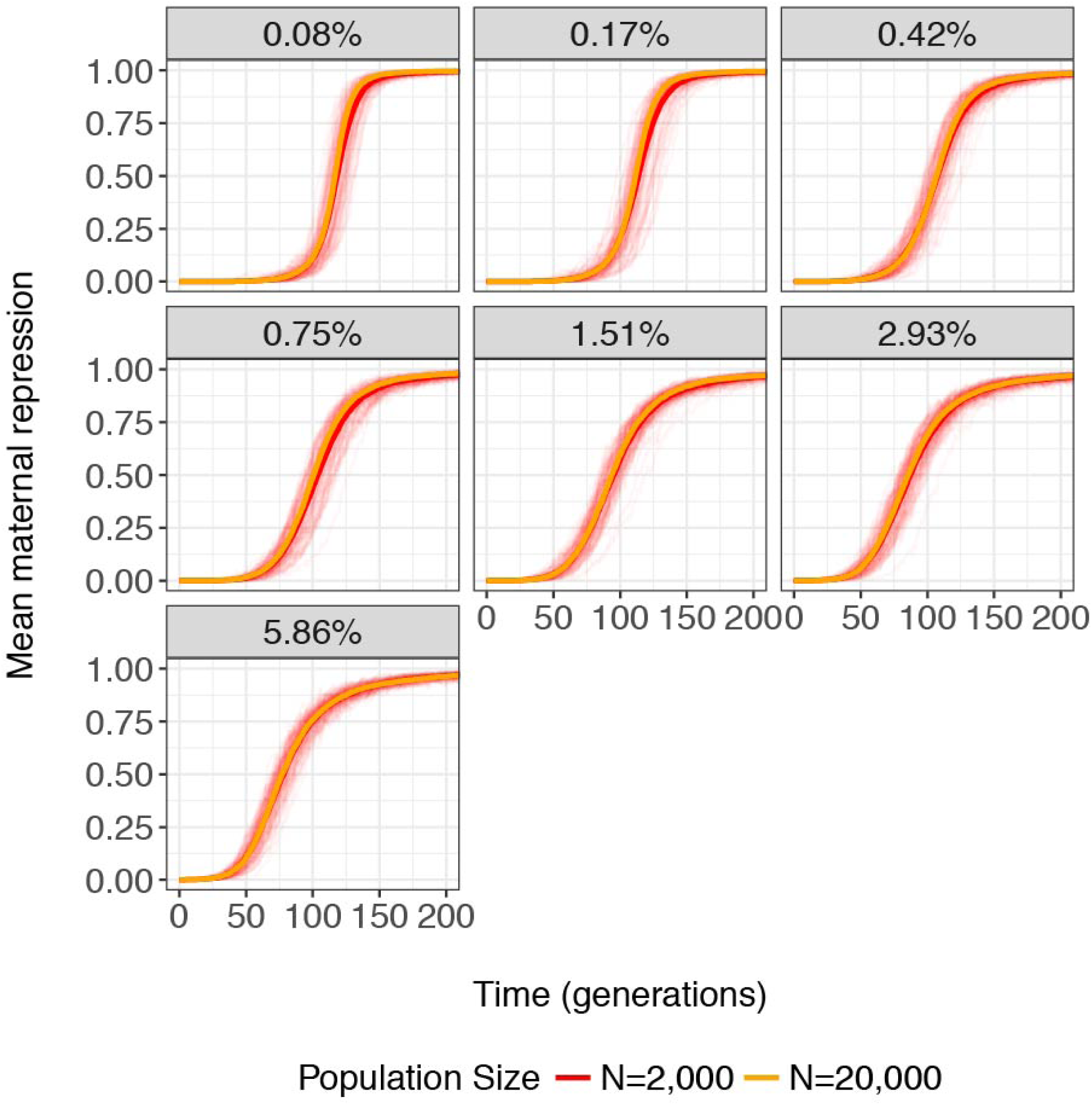
Larger populations do not exhibit repression earlier in invasion. The abundance of small-RNA sites (genomic percentage) is related to the evolution of mean maternal repression (*R*) over 200 generations in replicate simulated populations of *N* = 2,000 (red) and *N* = 20,000 (orange).The thick line represents the average response across 100 replicate populations for *N* = 2,000, 30 replicate populations for *N* = 20,000 and a small-RNA site percentage of 2.93%, and 10 replicate populations for all other small-RNA site percentages.

## Discussion

We harnessed the deep body of literature on *P*-element transposition and fitness effects (reviewed in Kelleher, 2016), as well as piRNA-mediated silencing of TEs (reviewed in Senti and Brennecke, 2010), in *D. melanogaster* to construct a forward simulation model of TE invasion and the evolution of host repression that is grounded in known biology. The validity of our model is supported by the degree to which our findings mirror empirical observations of *P*-element invasion in *D. melanogaster*. *P*-elements transitioned from rare to ubiquitous in natural population in *<* 20 years (Anxolabéhère et al., 1988), a rapid timescale that is matched by our simulated populations, in which invasion occurred in *<* 100 generations (figure 1A). Similarly, average *P*-element copy numbers in individual genomes in our simulated populations were in the range *∼* 5*−*60, which matches estimates from wild-derived genomes (Ronsseray et al. (1989); Srivastav and Kelleher (2017), figure 1D,E). Finally, repression in many natural populations evolved in *<* 30 years (Kidwell, 1983), which is echoed by our observation that the vast majority of populations evolve repression in *<* 200 generations (figure 1C). Although our model is, by design, informed by the specific biology of *P*-elements in *D. melanogaster*, our findings have general implications for TE invasion and the evolution of TE repression. Horizontal transfer of TEs and post-invasion TE proliferation are ubiquitous across the tree of life (reviewed in Schaack et al., 2010). Similarly, small-RNA-mediated silencing is a ubiquitous mechanism for TE control, employed by both prokaryotes and eukaryotes (reviewed in Blumenstiel, 2011). Our results therefore provide fundamental insights about the trajectory of TE invasion into new genomes, and the regulatory response of genomes to new invaders.

### Weak selection for repression of transposition?

Charlesworth and Langley (1986) suggested that selection for transpositional repression was likely to be weak in sexually reproducing organisms, because recombination would rapidly separate repressor alleles from chromosomes that they have protected from incurring deleterious mutations. Furthermore, they proposed that the adaptive evolution of transpositional repression relied on a sufficiently high frequency of dominant deleterious insertions and a sufficiently large effective population size. Consistent with this prediction, previous simulations of small-RNA-mediated retrotransposon silencing, which assumed dominant deleterious effects of new insertions and large effective population sizes, observed selection for repressor alleles (Lu and Clark, 2010). It is counter-intuitive, therefore, that we also observed positive selection for repressor alleles in relatively small populations of *N* = 2,000 individuals, with entirely recessive effects of TE insertions on fitness.

The apparent contradiction between our findings and the predictions of Charlesworth and Langley (1986) are at least partially explained by differences in the transposition rates we considered. The empirically estimated transposition rate for an unmarked *P*-element of 0.1 (new insertions/element/genome) (Eggleston et al., 1988) is multiple orders of magnitude higher than “typical” estimates of transposition rate in *Drosophila* (*∼* 10^*−*4^ *−*10^*−*5^; Nuzhdin, 1999). The high transposition rates of *P*-elements may be more realistic, however, because other transposition rate estimates were obtained from TE families we now understand to be regulated (*i.e.*, lowered) by the piRNA pathway (Brennecke et al., 2007; Nuzhdin, 1999). Higher transposition rates cause greater fitness benefits of repression, because they increase the likelihood of a deleterious insertion and therefore the relative advantage of an un-inserted chromosome.

The fitness model that we employed in this study, which was based on the transpositional bias of *P*-elements into functional sites (Bellen et al., 2011, 2004; Spradling et al., 2011), and the estimated fitness effects of individual *P*-element insertions (Cooley et al., 1988; Mackay et al., 1992), likely also contributes to the strength of selection we observe. Although they act recessively, the empirically estimated fitness effects of *P*-element insertions are quite deleterious, with an average fitness cost of 12.2%, and 3.7% of sites acting as recessive lethals (Cooley et al., 1988; Lyman et al., 1996; Mackay et al., 1992). Similar to high transposition rates, these dramatic fitness consequences enhance the benefits of repressing transposition. Transposable elements exhibit a diversity of preferences for particular insertion sites, and the degree to which these preferences influence selection for transpositional repression remains unknown.

### Repression of dysgenic sterility and ectopic recombination enhances adaptive evolution of small-RNA-mediated TE silencing

In addition to deleterious insertions, we also revealed that dysgenic sterility and ectopic recombination can exert positive selection on repressor alleles in simulated populations (figure 3C, 5B). Our results differ from those of Lee and Langley (2012), who previously considered selection on small-RNA-independent repressors of dysgenic sterility. They observed that strong selection on the repressor allele was too ephemeral to produce significant changes in its population frequency, owing to the rapid loss of non-repressive maternal genotypes from the population. However, they also assumed that the presence of *P*-elements in the maternal genome was sufficient to establish small RNA silencing, rather than explicitly modeling the production of regulatory small RNAs from insertions at particular sites as we do here. Our observations therefore suggest that dysgenic sterility can be an important agent of selection for TE regulation, at least during TE invasion. Dysgenic sterility may further be a common cost of TEs that selects for repression in their hosts. In *Drosophila*, multiple historic TE invasions are associated with dysgenic sterility syndromes (Bingham et al., 1982; Blackman et al., 1987; Bucheton et al., 1984; Evgen’ev et al., 1997; Hill et al., 2016), and at least three of these syndromes have been attributed to the absence of maternally deposited piRNAs (Blumenstiel and Hartl, 2005; Brennecke et al., 2008; Chambeyron et al., 2008; Rozhkov et al., 2010). More broadly, mutations that impact TE regulation are associated with fertility defects in diverse eukaryotes (reviewed in Castañeda et al., 2011; Mani and Juliano, 2013).

In forward simulations of TE dynamics, costs of ectopic recombination are described by a fitness model that assumes negative epistasis among TE insertions (Dolgin and Charlesworth, 2006, 2008; Lee and Langley, 2012; Lu and Clark, 2010). Negative epistasis appears to be required to stabilize genomic TE copy numbers in the absence of repression (Charlesworth and Charlesworth, 1983). However, small-RNA-mediated silencing may reduce the fitness costs of ectopic recombination by establishing a heterochromatic environment at TE loci that suppresses recombination (reviewed in Blumenstiel, 2011). Therefore, ectopic recombination could also select for small-RNA-mediated repression. Our observations supports this prediction. We observed enhanced positive selection on small-RNA-mediated repressors at higher ectopic recombination rates (figure 4B, C). Because TE copy numbers remained relatively low in the simulations we presented here (generally *<* 60 copies), positive selection could be much stronger in large, repeat-rich genomes, where individual elements can be represented by thousands of copies (*e.g.*, Naito et al., 2006). Suppression of ectopic recombination may therefore represent a major benefit of small-RNA-mediated repressor alleles.

In contrast to the benefits of repressing transposition and potentially ectopic recombination, heterochromatin formation at TE loci (*i.e.*, transcriptional silencing) can also have a deleterious effect if it interferes with host gene expression. Such deleterious effects have been observed in both *Arabidopsis* (Hollister and Gaut, 2009) and *Drosophila* (Lee, 2015). Exploring trade-offs between the costs and benefits of repression, and how such trade-offs influence transcriptional versus post-transcriptional regulation of TEs, would make an interesting direction of future study.

### The abundance of small-RNA-producing sites is a major determinant of post-invasion dynamics

In the *D. melanogaster* genome we used as a model for our simulations the small RNAs that regulated TEs are thought to be encoded by *>* 100 independent loci, comprising *∼*3% of genomic sites(Brennecke et al., 2007). Small RNA mediated silencing is therefore a polygenic trait encoded by a large mutational target, a property that is shared with other taxa. Although the power to identify loci that give rise to TE-regulating small RNAs can be limited by the quality of the genome assembly as well as its repeat content, large numbers of repeat-rich small-RNA-producing sites are also implicated in TE regulation in mouse (Aravin et al., 2006), *Caenorhabditis* (Ni et al., 2014) and even *Arabidopsis* (reviewed in Lisch and Bennetzen, 2011).

Large mutational target sizes are predicted to result in adaptation via soft selective sweeps (Pritchard et al., 2010), in which multiple beneficial alleles increase in frequency but do not reach fixation (Pennings and Hermisson 2006a,b). Throughout our simulations, we observed exactly these soft-sweeps, in which beneficial insertions in small-RNA-producing sites segregate at higher frequency than their neutral counterparts, but rarely reach fixation (Figure 2B). Our results are consistent with the plethora of independent P-element insertions into piRNA clusters that have been reported in natural populations of *D. melanogaster* (Marin et al. 2000; Ronsseray et al. 1996; Stuart et al. 2002, reviewed in Kelleher 2016). We furthermore discovered that that the larger the mutational target (*i.e.* the more common small-RNA-producing sites are in the genome), the weaker selection for repression becomes, because the mutational load of deleterious insertions that select for repression is decreased (Figure 5A, D, Supplementary Figure 5). To our knowledge, this unexpected relationship between mutational target size and positive selection is unique to the evolution of TE repression, since the enhanced benefits of repression when the mutational target is small reflect accumulating TE copy numbers.

Why are small-RNA producing sites so common in host genomes? An appealing adaptive explanation is that the large mutation target facilitates the establishment of silencing when new TE families invade (Kelleher, 2016). However, our findings do not support this model, as repression evolved within *∼* 200 generations regardless of the number of small-RNA sites in the theoretical genome (figure 5A). Rather, we observed that the mutational target size has a dramatic impact on the maximum copy number the TE achieves after invasion, with genomes containing fewer small-RNA sites accumulating many more TE copies (figure 5C). This accumulation of TE copies can be attributed to longer individual waiting times for the occurrence of the first repressor allele, during which time TEs enjoy unrestricted transposition (figure 5A, B, D).

The magnitude of interspecific variation in the proportion of the genome that corresponds to small-RNA-producing sites remains largely unknown. In part this reflects the challenge of annotating such sites, which lack diagnostic primary sequence features, and whose transcripts are rapidly processed into short sequences that align to multiple genomic locations (Brennecke et al., 2007; Rosenkranz and Zischler, 2012). Nevertheless, our observations suggest an unexpected benefit to having more small-RNA sites, namely, that the larger mutational target buffers genomes against post-invasion proliferation. They further provide a potential explanation for rapid, TE-mediated expansions in genome size, which have been observed in certain vertebrate and plant lineages (Lee and Kim, 2014; Sun and Mueller, 2014). If the relative fraction of the genome that corresponds to small-RNA-producing sites decreases when genome size expands, a type of snowballing could occur, in which large genomes experience increasingly dramatic TE proliferations and corresponding decreases in the relative abundance of small-RNA-producing sites.

## Materials and Methods

### Statistical Analyses

All statistical analyses were conducted with R version 3.3 (R Core Team, 2016). All graphs were produced in ggplot2 (Wickham, 2009).

### Data Availability

The software used to run all simulations was written in Perl and will be available at https://github.com/… at the time of publication.

## Supplementary Material

Supplementary table S1 and figures S1–S4 are available at Genome Biology and Evolution online (https://academic.oup.com/gbe/).

## Acknowledgments

R. Meisel and four anonymous reviewers provided useful comments on the manuscript. We used the Maxwell cluster from the Center of Advanced Computing and Data Systems (CACDS) at the University of Houston. CACDS staff provided technical support. The National Science Foundation (grant DEB-1457800 awarded to E.S.K. and grant DEB-1354952 awarded to R.B.R.A.) funded this work.

